# Anthracyclines inhibit SARS-CoV-2 infection

**DOI:** 10.1101/2023.01.10.523518

**Authors:** Zhen Wang, Qinghua Pan, Ling Ma, Jianyuan Zhao, Fiona McIntosh, Zhenlong Liu, Shilei Ding, Rongtuan Lin, Shan Chen, Andrés Finzi, Chen Liang

**Author notes:** Address correspondence to Chen Liang.

## Abstract

Vaccines and drugs are two effective medical interventions to mitigate SARS-CoV-2 infection. Three SARS-CoV-2 inhibitors, remdesivir, paxlovid, and molnupiravir, have been approved for treating COVID-19 patients, but more are needed, because each drug has its limitation of usage and SARS-CoV-2 constantly develops drug resistance mutations. In addition, SARS-CoV-2 drugs have the potential to be repurposed to inhibit new human coronaviruses, thus help to prepare for future coronavirus outbreaks. We have screened a library of microbial metabolites to discover new SARS-CoV-2 inhibitors. To facilitate this screening effort, we generated a recombinant SARS-CoV-2 Delta variant carrying the nano luciferase as a reporter for measuring viral infection. Six compounds were found to inhibit SARS-CoV-2 at the half maximal inhibitory concentration (IC50) below 1 μM, including the anthracycline drug aclarubicin that markedly reduced viral RNA-dependent RNA polymerase (RdRp)-mediated gene expression, whereas other anthracyclines inhibited SARS-CoV-2 by activating the expression of interferon and antiviral genes. As the most commonly prescribed anti-cancer drugs, anthracyclines hold the promise of becoming new SARS-CoV-2 inhibitors.

**IMPORTANCE:** Microbial metabolites are a rich source of bioactive molecules. The best examples are antibiotics and immunosuppressants that have transformed the practice of modern medicine and saved millions of lives. Recently, some microbial metabolites were reported to have antiviral activity, including the inhibition of Zika virus and Ebola virus. In this study, we discovered several microbial metabolites that effectively inhibit SARS-CoV-2 infection, including anthracyclines that have also been shown to inhibit other viruses including Ebola virus through enhancing interferon responses, which indicates potentially broad antiviral properties of these microbial metabolites and can lead to the discovery of pan-antiviral molecules.

## INTRODUCTION

Coronavirus disease 2019 (COVID-19) is caused by severe acute respiratory syndrome coronavirus 2 (SARS-CoV-2) (1, 2). Since World Health Organization (WHO) declared COVID-19 pandemic on March 11, 2020, there have been 656 million confirmed COVID-19 cases and over 6.7 million deaths. The impacts of COVID-19 on public health, social life and economy have been astronomical. In addition to timely diagnosis, quarantine, social distancing and masking, COVID vaccines have played a critical role in preventing SARS-CoV-2 infection, reducing disease severity and fatality, and restoring normal life (3). For those who contract SARS-CoV-2, effective treatments become pivotal in preventing disease progression and saving lives (4).

At the beginning of COVID-19 pandemic, treatment options were limited to supportive care such as oxygen and mechanical ventilation, immunomodulatory therapies with dexamethasone and adjunctive therapies with anticoagulants (4). A series of clinical trials were quickly conducted to test repurposed drugs including hydrocloroquine and ivermectin, but no significant clinical benefits were observed (5–8). Inconsistent results were reported from various trials of convalescent plasma, likely owing to unselected patients, including the vaccination status (9). Neutralizing monoclonal antibodies such as casirivimab plus imdevimab, bamlanivimab plus etesevimab, sotrovimab, tixagevimab plus cilgavimab have shown clinical benefits for both in-patients and out-patients (10, 11), and have been approved to treat COVID-19 patients with mild to moderate symptoms. Remdesivir was the first direct antiviral drug approved to treat COVID-19 (12, 13), followed by molnupiravir (14) and paxlovid (15) which are used to treat patients at the onset of symptoms to suppress viral replication. The constantly emerging SARS-CoV-2 variants, in particular the Omicron variants bearing more than 50 mutations with over 30 in the viral Spike glycoprotein, often defy the effectiveness of vaccines and antibody-based therapy and pose resistance to the few approved antiviral drugs (16–18).

With this challenge, considerable efforts have been made to discover new, effective anti-SARS-CoV-2 compounds, which can also serve as a reserve to repurpose and treat future emerging human coronaviruses. Along this line, we screened a library of 620 microbial metabolites for compounds with anti-SARS-CoV-2 activity. We chose to examine microbial metabolites because they have been a rich source of bioactive molecules in modern medicine (19). The prominent examples include antibiotics, anti-cancer drugs, and immunosuppressants which dramatically increase the success of organ transplantation (19). Recently, microbial metabolites with promising antiviral activity have been reported to inhibit Zika virus (20), influenza virus (21), and others (22), and may serve as useful source for the discovery of antiviral inhibitors including against SARS-CoV-2. To facilitate the screening effort, we used nano luciferase (NLuc) to replace the ORF7a gene in SARS-CoV-2 Delta variant genome, and produced the SARS-CoV-2 Delta NLuc reporter virus. We have successfully identified 6 compounds inhibiting SARS-CoV-2 infection of the lung epithelial cell line Calu-3, with aclarubicin and ecteinascidin-770 showing the most potent inhibitory activity.

## MATERIALS AND METHODS

### Cells and viruses

Calu-3 (cat. HTB-55), Vero E6 (cat. CRL-1586), HEK293 (cat. CRL-1573) and HEK293T (cat. CRL-3216) cells were purchased from ATCC. Calu-3 cells were grown in Eagle’s minimum essential medium (EMEM) (Wisent, cat. 320-005-CL) supplemented with 15% fetal bovine serum (FBS) (Gibco, cat. 12483-020). Vero E6 and HEK293T cells were grown in Dulbecco’s modified Eagle medium (DMEM) supplemented with 10% FBS. Vero E6/TMPRSS2 cell line was generated by transducing Vero E6 cells with lentiviral particles expressing human TMPRSS2, followed by selection with 1 mg/ml G418 (Wisent, cat. 400-130-IG). HEK293/ACE2/TMPRSS2 cells were generated by sequential transduction of HEK293 cells with lentiviral particles expressing human ACE2 (selection with 2 μg/ml puromycin) and lentiviral particles expressing human TMPRSS2 (selection with 1 mg/ml G418). The TMPRSS2 (cat. 145843) and ACE2 (cat. 145839) lentiviral DNA clones (23) were purchased from Addgene and transfected into HEK293T cells together with pSPAX2 (Addgene, cat. 12260) and VSV-G DNA (Addgene, cat. 8454) (24) to produce lentiviral particles. Virus HCoV-229E-Luc was kindly provided by Volker Thiel (25). Sendai virus (Cantel Strain) was obtained from Charles River Laboratories (26).

### Compounds

The microbial metabolite library (HY-L084) was purchased from MedChemExpress. Aclarubicin hydrochloride (cat. HY-N2306A, CAS No. 75443-99-1), daunorubicin hydrochloride (cat. HY-13062, CAS No. 23541-50-6), doxorubicin hydrochloride (cat. HY-15142, CAS No. 25316-40-9), epirubicin hydrochloride (cat. HY-13624A, CAS No. 56390-09-1), idarubicin hydrochloride (cat. HY-17381, CAS No. 57852-57-0), mitoxantrone dihydrochloride (cat. HY-13502A, CAS No. 70476-82-3), and ecteinascidin 770 (cat. HY-101191, CAS No. 114899-80-8) were purchased from MedChemExpress. Remdesivir (cat. 30354, CAS No. 1809249-37-3) was purchased from Cayman Chemical, nirmatrelvir (cat. T9351, CAS No. 2628280-40-8) from Targetmol.

### Generation of the SARS-CoV-2 Delta NLuc BAC DNA clone

The complete SARS-CoV-2 Delta variant sequence (Genbank accession number MZ724531.1) was used to construct the infectious SARS-CoV-2 DNA clone. The nano luciferase (NLuc) gene was used to replace the ORF7a as previously described (27). The CMV early promoter, HDV ribozyme sequence (Rz) and bovine growth hormone mRNA polyA signal sequence (bGH), as described in (28), were added to the 5’ and 3’ ends of Delta variant sequence to allow transcription and proper processing of the 3’ end of SARS-CoV-2 RNA (Fig. 1A). The SARS-CoV-2/NLuc DNA was synthesized by DNA Codex Inc, cloned into the BAC (bacterial artificial chromosome) vector, transformed into TransforMax EPI300 electrocompetent cells (Lucigen) by electroporation, and prepared with ZR BAC DNA Miniprep Kit (ZYMO RESEARCH, cat. D4048). The sequence of SARS-CoV-2 Delta NLuc BAC DNA was verified by next-generation sequencing (NGS).

**Figure 1.**
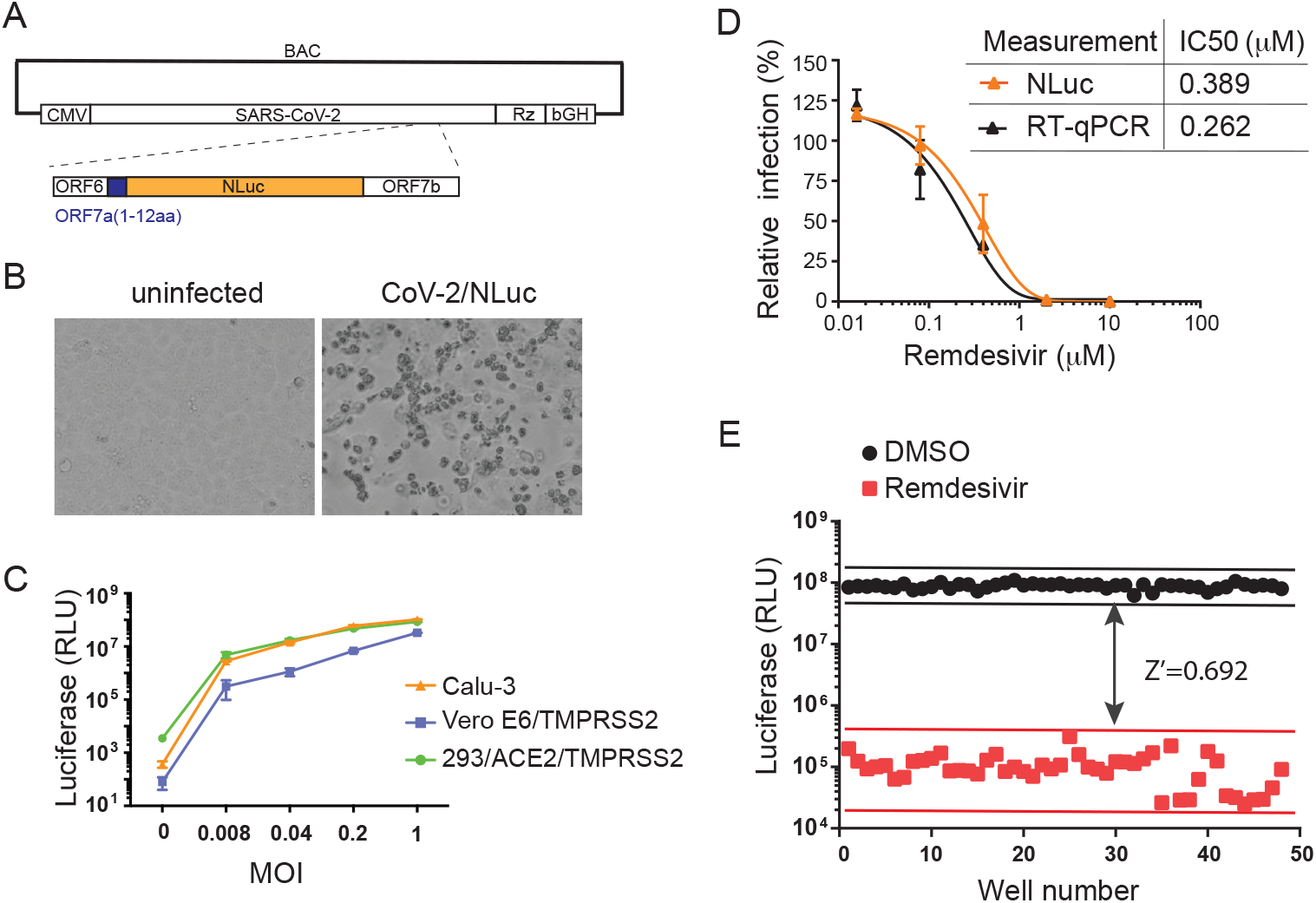
Production of a recombinant SARS-CoV-2 Delta/NLuc virus. (A) Depiction of the BAC vector containing the SARS-CoV-2 Delta DNA. The NLuc sequence replaces viral ORF7a gene. The first 12 amino acids of ORF7a are retained. (B) Vero E6/TMPRSS2 cells were infected with CoV-2/NLuc. The recorded cytopathogenic effect was at day 5 after infection. (C) NLuc activity was measured in Calu-3, Vero E6/TMPRSS2, and 293/ACE2/TMPRSS2 cells that were infected with CoV-2/NLuc of different MOIs for 24 hours. (D) Different concentrations of remdesivir was added to Calu-3 cell culture when infected with CoV-2/NLuc (MOI=0.2). Forty hours after infection, NLuc activity in the infected cells was measured. Levels of viral RNA in culture supernatants were determined by RT-qPCR. Each remdesivir concentration was tested in triplicate infections. IC50 of remdesivir inhibition was calculated using the NLuc and viral RNA data. (E) CoV-2/NLuc (MOI=0.2) was used to infect Calu-3 cells seeded in a 96-well plate. Forty-eight wells were treated with 10 μM remdesivir, the other 48 wells were exposed to 1% DMSO. NLuc activity was measured 40 hours after infection. Z score was calculated as described in a previous study (38).

### Production of SARS-CoV-2

SARS-CoV-2 RIM-1, SA and BA.1 strains were generously provided the MUHC (McGill University Health Centre) clinical microbiology lab and LSPQ (Laboratoire de santé publique du Québec). The RIM-1 strain was isolated from a patient at MUHC (GenBank accession number MW599736; lineage B.1.147). SARS-CoV-2 was amplified by infecting Vero E6 cells in DMEM containing 2% FBS. Viruses were harvested when over 70% of cells exhibited cytopathic effect (CPE), by centrifugation at 1500 g for 10 min and filtration through a 0.2 μm filter to remove cell debris, then aliquoted and stored at −80°C. The titer of virus stock was determined with the TCID50 (median tissue culture infectious dose) assay. Briefly, viral dilutants (1:10) were used to infect Vero E6 cells in a 48-well plate for 4 days before cells were fixed with 10% formaldehyde for 1 hour, followed by staining with 0.1% crystal violet (in 10% ethanol) for 30 min. Wells with CPE were scored, and TCID50 values were calculated using the Reed&Muench method. To produce the CoV-2/NLuc virus, BAC DNA was transfected into HEK293/ACE2/TMPRSS2 cells using lipofectamine 3000 (Thermo Fisher, cat. L3000001). Two days after transfection, supernatants were used to infect Vero E6/TMPRSS2 cells to amplify the virus. At passage 3, viral RNA was extracted from culture supernatants and sequenced by NGS. The amplified virus presented the same sequence as the synthesized CoV-2/NLuc. Titres of viruses were determined by TCID50 assays. SARS-CoV-2 infection was performed in the biocontainment level 3 facility at the Research Institute of MUHC, approved by the Public Health Agency of Canada.

### Testing compounds against SARS-CoV-2

Inhibition of CoV-2/NLuc infection was evaluated in 96-well plates. Calu-3 cells were seeded in a black wall 96-well plate (Greiner Bio-One, cat. 655086) at 2×10^4^ cells/well. Two days after, cells were exposed to CoV-2/NLuc (MOI=0.2) for 1 hour in the presence of the compound. Control infection was performed with the solvent (such as 1% dimethyl sulfoxide (DMSO)) of the compound. Viruses were then washed off, cells were cultured together with the compound for 40 hours before luciferase activity in the infected cells was measured using Nano-Glo Luciferase Assay System (Promega, cat. N1120). Inhibition of SARS-CoV-2 variants RIM-1, SA and BA.1 was tested in 24-well plates. Calu-3 cells were seeded at 1.5×10^5^ cells per well two days before the infection started for 1 hour in the presence of the compound. After viruses were washed off, cells were cultured together with the compound for 40 hours before culture supernatants were collected to extract viral RNA with TRIzol LS (Thermo Fisher, cat. 10296010). Levels of viral RNA were determined by RT-PCR using a primer pair (nCoV_IP2-12669Fw: ATGAGCTTAGTCCTGTTG, nCoV_IP2-12759Rv: CTCCCTTTGTTGTGTTGT, probe: AGATGTCTTGTGCTGCCGGTA [5’]Fam [3’]BHQ-1) that amplifies a 108 bp fragment of viral RdRp gene. RT-PCR was performed with TaqMan Fast Virus 1-Step Master Mix (Invitrogen, cat. 4444434) in QuantStudio 7 Flex Real-Time PCR System (Thermo Fisher). The relative change of viral RNA between the compound-treated infection samples and the control infection samples was calculated using the ΔCt values. Each compound concentration was tested in triplicate infections.

### Cytotoxicity assay

Calu-3 cells were seeded in a white wall 96-well plate (Greiner Bio-One, cat.655074) at 2×10^4^ cells/well. Two days after, cells were treated with compounds and cultured for 40 hours. Then, 100 μl CellTiter-Glo Reagent (CellTiter-Glo Luminescent Cell Viability Assay, Promega, cat. G7571) was added to each well for 10 min before luminescence was measured with PerkinElmer EnSight microplate reader. Each compound concentration was tested in triplicate.

### Cell-based SARS-CoV-2 RdRp assay

The assay was performed as we previously described (29). Briefly, HEK293T cells were transfected with pCoV-Gluc plasmid DNA either alone or with pCOVID19-nsp12, pCOVID19-nsp7, pCOVID19-nsp8 at a plasmid ratio of 1:10:30:30. Twelve hours after transfection, cells were seeded into a 96-well plate at 10^4^ cells/well, followed by treatment with various doses of aclarubicin or ecteinascidin 770 for 24 hours. Each drug concentration was tested in triplicate. 10 μl of the supernatant was transferred into a white wall 96-well plate, and 60 μl of 16.7 μM coelenterazine-h was injected to measure luminescence using the Berthold Centro XS3 LB 960 microplate luminometer.

### Cell-based SARS-CoV-2 protease assay

The EGFP-RLuc fusion protein has the ITSAVLQSGFRK sequence inserted between EGFP and RLuc protein, which serves as the cleavage site of SARS-CoV-2 main protease 3CLPro (30). The EGFP-RLuc plasmid (100 ng) was transfected into HEK293T cells in a 6-well plate alone or with the 3CLPro plasmid (300 ng). Twelve hours after transfection, cells were treated with nirmatrelvir (1 μM), aclarubicin (800 nM), ecteinascidin 770 (10 nM), or DMSO (1%) for 48 hours before cells were harvested for Western blotting to detect the uncleaved EGFP/RLuc and the cleaved EGFP using anti-GFP antibodies (Santa Cruz, cat. sc-9996).

### Measuring interferon production and ISG expression

Levels of IFN-λ in the culture supernatants were determined with ELISA using the human IL-29 ELISA kit (Invitrogen, cat. BMS2049) following the instructions of the manufacturers. Expression of MxB, IFITM3 and ISG15 in the treated cells was examined by Western blotting using antibodies against MxB (31), IFITM3 (Proteintech, cat.11714-1-AP), or ISG15 (Abcam, cat. ab48020). Tubulin was detected as the control. Briefly, cells were lysed in RIPA buffer (50 mM Tris-HCl [pH 7.5], 150 mM NaCl, 1 mM EDTA, 1% Nonidet P-40, 0.1% SDS) containing 1xprotease inhibitor cocktail (Roche) on ice for 30 min before centrifugation in an Eppendorf microcentrifuge at 4°C. Cell lysates of 20 μg protein were separated in a SDS-12% polyacrylamide gel (PAGE) followed by transferring proteins to a polyvinylidene difluoride (PVDF) membrane (Roche, catalog 3110040). After blocking in 5% milk (in 1xphosphate buffered saline (PBS)) at room temperature for 1 hour, the membrane was incubated with the indicated primary antibodies at room temperature for 2 hours, washed with 1xPBS containing 0.1% Tween 20, then incubated with secondary antibodies conjugated with horseradish peroxidase (HRP). The membranes were treated with chemiluminescence reagents (PerkinElmer) before exposure to X-ray films.

### ISRE-Luc reporter assay

The ISRE-Luc reporter plasmid (32) (25 ng) was either transfected alone or co-transfected with a plasmid expressing SARS-CoV-2 genes NSP13 (Addgene, cat. 141379) (25 ng), ORF3a (Addgene, cat.141383) (25 ng) or ORF6 (Addgene, cat. 141387) (25 ng) (33), into HEK293T cells in a white wall 96-well plate. The transfected cells were treated with anthracyclines or Sendai virus (40 hemagglutinin [HA] units/ml) for 24 hours before luciferase activity was determined with the Steady-Glo luciferase assay system (Promega, cat. E2520).

### Statistical analysis

All statistical analyses were performed with GraphPad Prism version 9 software. Statistics were calculated using an unpaired t-test (two-tailed). Data shown are means of three independent experiments ± SD. Statistically significant differences are shown as *p<0.05, **p<0.01, ***p<0.001.

## RESULTS

### Production of SARS-CoV-2 Delta NLuc reporter virus

Several reverse genetics approaches have been reported by different groups to produce recombinant SARS-CoV-2 (34). One key step of these different approaches is to generate the full-length 30 kb SARS-CoV-2 DNA by assembling viral DNA fragments either *in vitro* through the use of restriction enzymes (27) and circular polymerase extension reaction (CPER) (35), or in yeast by homology recombination (36). The full-length viral RNA is then *in vitro* synthesized using T7 RNA polymerase from the assembled viral DNA with the T7 promoter attached to its 5’ end, followed by transfection into cells to produce live SARS-CoV-2. Alternatively, the cytomegalovirus (CMV) early promoter is attached to the 5’ end of SARS-CoV-2 DNA and drives transcription when this DNA clone is transfected into cells (36). To generate the correct 3’ end of viral RNA, the hepatitis D virus (HDV) ribozyme is appended to the 3’ end of SARS-CoV-2 DNA followed by the bovine growth hormone poly(A) signal (37). To produce live recombinant SARS-CoV-2 containing a luciferase reporter gene, we have the SARS-CoV-2 Delta variant genome DNA synthesized with viral ORF7a gene replaced with nano luciferase (NLuc) sequence. The CMV promoter was attached to the 5’ end, the HDV ribozyme to the 3’end of viral DNA, and this viral DNA was cloned into the bacterial artificial chromosome (BAC) vector (Fig. 1A). This CoV-2/NLuc DNA clone was transfected into HEK293/ACE2/TMPRSS2 cells followed by propagation in Vero E6/TMPRSS2 cells. Cytopathic effects were observed (Fig. 1B), indicating successful viral replication. At the 3^rd^ passage, viral RNA from 5 independent viral cultures was extracted and sequenced. The results of next-generation sequencing (NGS) confirmed that 2 viral clones had the same sequence as the synthetic CoV-2/NLuc DNA. Titer of the virus stock was determined in TCID50 (median tissue culture infectious dose) assay and equated to 3.18×10^6^ PFU/ml. We then infected Calu-3 cells with CoV-2/NLuc of increasing MOIs, and detected increasing levels of NLuc in the infected cells (Fig. 1C). To demonstrate the utility of CoV-2/NLuc reporter virus in testing anti-CoV-2 compounds by measuring NLuc, we examined increasing doses of remdesivir, a viral RNA-dependent RNA polymerase (RdRp) inhibitor, in inhibiting CoV-2/NLuc infection of Calu-3 cells. CoV-2/NLuc infection was monitored either by measuring NLuc in the infected Calu-3 cells or by quantifying viral RNA in the culture supernatants by quantitative reverse transcription (RT)-PCR. Similar remdesivir inhibition curves were generated with these two methods of quantifying CoV-2/NLuc infection, with IC50=0.389 μM by measuring NLuc and 0.262 μM by measuring viral RNA (Fig. 1D). To determine whether CoV-2/NLuc is suitable for discovering antiviral compounds, we determined the Z score which reports the statistical effect size and signal window of an assay, by measuring NLuc in CoV-2/Nluc infected Calu-3 cells that were treated with or without 10 μM remdesivir. The Z score was calculated as 0.692 (Fig. 1E), satisfying the minimum requirement of Z=0.5 (38). We have therefore produced recombinant CoV-2/NLuc virus whose infection can be quantified by measuring NLuc in the infected cells and can be used to screen for antiviral compounds.

### Identification of microbial metabolites with anti-SARS-CoV-2 activity

The microbial metabolite library from MedChemExpress has a collection of 620 compounds from bacteria and fungi. A variety of biological activities and functions have been reported for compounds in this library, including antiviral activity (Fig. 2A). Our aim was to determine whether any of these microbial metabolites inhibits CoV-2/NLuc replication in Calu-3 cells. The fold of decrease in NLuc in the infected cells upon exposure to compounds reports the degree of inhibition of CoV-2/NLuc (Fig. 2A). The initial test identified 55 compounds exhibiting more than 5-fold inhibition of CoV-2/NLuc at 10 μM concentration (Fig. 2B). To examine whether the decreased luciferase activity is a result of cytotoxicity caused by the compounds, viability of Calu-3 cells in the presence of 10 μM of each of these 55 compounds was measured. 70% or higher cell viability was observed for 24 compounds (Fig. 2C). We then tested serial concentrations of these compounds against CoV-2/NLuc infection of Calu-3 cells to determine their IC50 values. Viability of uninfected Calu-3 in the presence of these compounds was also determined. Six compounds presented an IC50 value lower than 1 μM (Fig. 3A, Table 1). These 6 compounds are aclacinomycin A hydrochloride (also called aclarubicin), anisomycin, ecteinascidin 770 (ET-770), geldanamycin, mycophenolic acid, and T-2 toxin. We further tested these 6 compounds for inhibiting the common cold human coronavirus HCoV-229E-Luc, which has the Renilla luciferase gene inserted in viral ORF4 (25). Anisomycin and T-2 toxin exhibited similar inhibition of both CoV-2/NLuc and HCoV-229E-Luc, whereas aclarubicin, ET-770, geldanamycin and mycophenolic acid showed much weaker inhibition of HCoV-229E-Luc (Fig. 3B, Table 1). We next tested the anti-CoV-2 activity of aclarubicin and ET-770 in HEK293T/ACE2/TMPRSS2 cells and observed strong inhibition of CoV-2/NLuc by both compounds (Fig. 3C). When SARS-CoV-2 variants RIM-1 (alpha variant), SA (beta variant) and BA.1 (omicron variant) were tested by measuring virus production in the culture supernatants with RT-qPCR, potent inhibition by both compounds was observed (Fig. 3D). Therefore, aclarubicin and ET-770 showed specific inhibition of SARS-CoV-2 in a cell-type independent manner.

**Figure 2.**
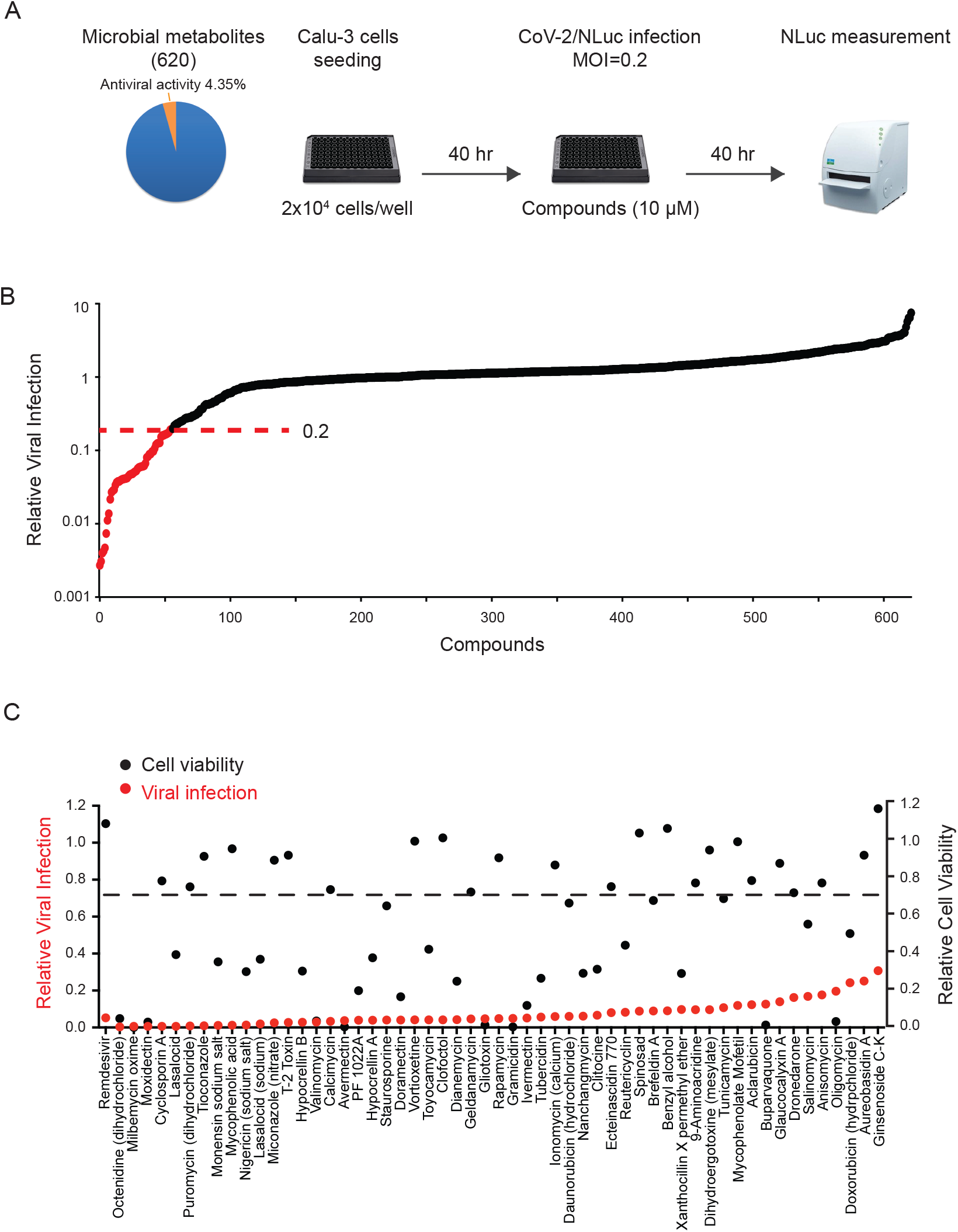
Identification of anti-SARS-CoV-2 microbial metabolites. (A) Testing the library of microbial metabolites for compounds inhibiting CoV-2/NLuc infection of Calu-3 cells (MOI=0.2). (B) Relative viral infection under treatment by each compound was calculated using the NLuc values of control infection (1% DMSO) arbitrarily set as 1. Shown in red are compounds that suppressed CoV-2/NLuc by more than 5 folds. Remdesivir was tested as a positive control. (C) Cytotoxicity of compounds at 10 μM in Calu-3 cells. Also shown are their folds of inhibition of CoV-2/NLuc infection (in red dots).

**Figure 3.**
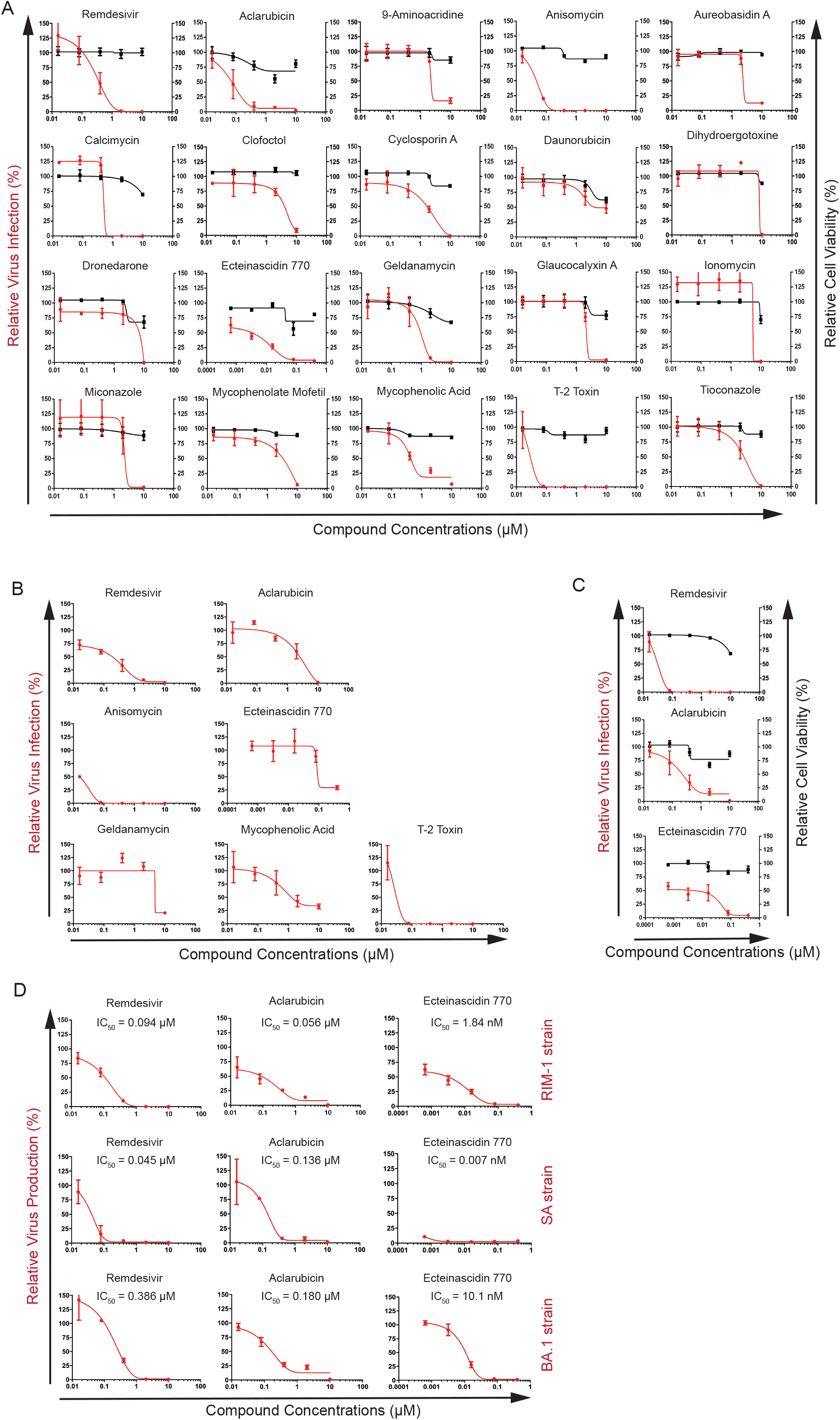
Determination of IC50 values of lead compounds against SARS-CoV-2 and hCoV-229E. (A) Serial dilutions of each lead compound were tested for inhibiting CoV-2/NLuc infection of Calu-3 cells (MOI=0.2). Each drug concentration was tested in triplicate infections. Also shown for each compound is cytotoxicity at the drug concentrations tested. Luciferase values of DMSO (1%) control are arbitrarily set as 100. (B) Inhibition of hCoV-229E/RLuc infection of Calu-3 cells (MOI=0.1) by the lead compounds. (C) Inhibition of CoV-2/NLuc infection of 293/ACE2/TMPRSS2 cells by aclarubicin and ecteinascidin 770. Cytotoxicity of each compound at the tested concentrations in 293/ACE2/TMPRSS2 cells was determined. Remdesivir was tested for inhibiting CoV-2/NLuc and hCoV-229E/RLuc as a positive control. (D) Aclarubicin and ecteinascidin 770 inhibit SARS-CoV-2 strains RIM-1, SA and BA.1. Serial dilutions of aclarubicin and ecteinascidin 770 were used to treat Calu-3 cells that were infected with either the RIM-1, SA or the BA.1 strain. Virus production in culture supernatants was determined by RT-qPCR to measure viral RNA levels. Viral RNA levels in control infection (1% DMSO) are arbitrarily set as 100. All drug concentrations were tested in triplicate infections.

**Table 1.**
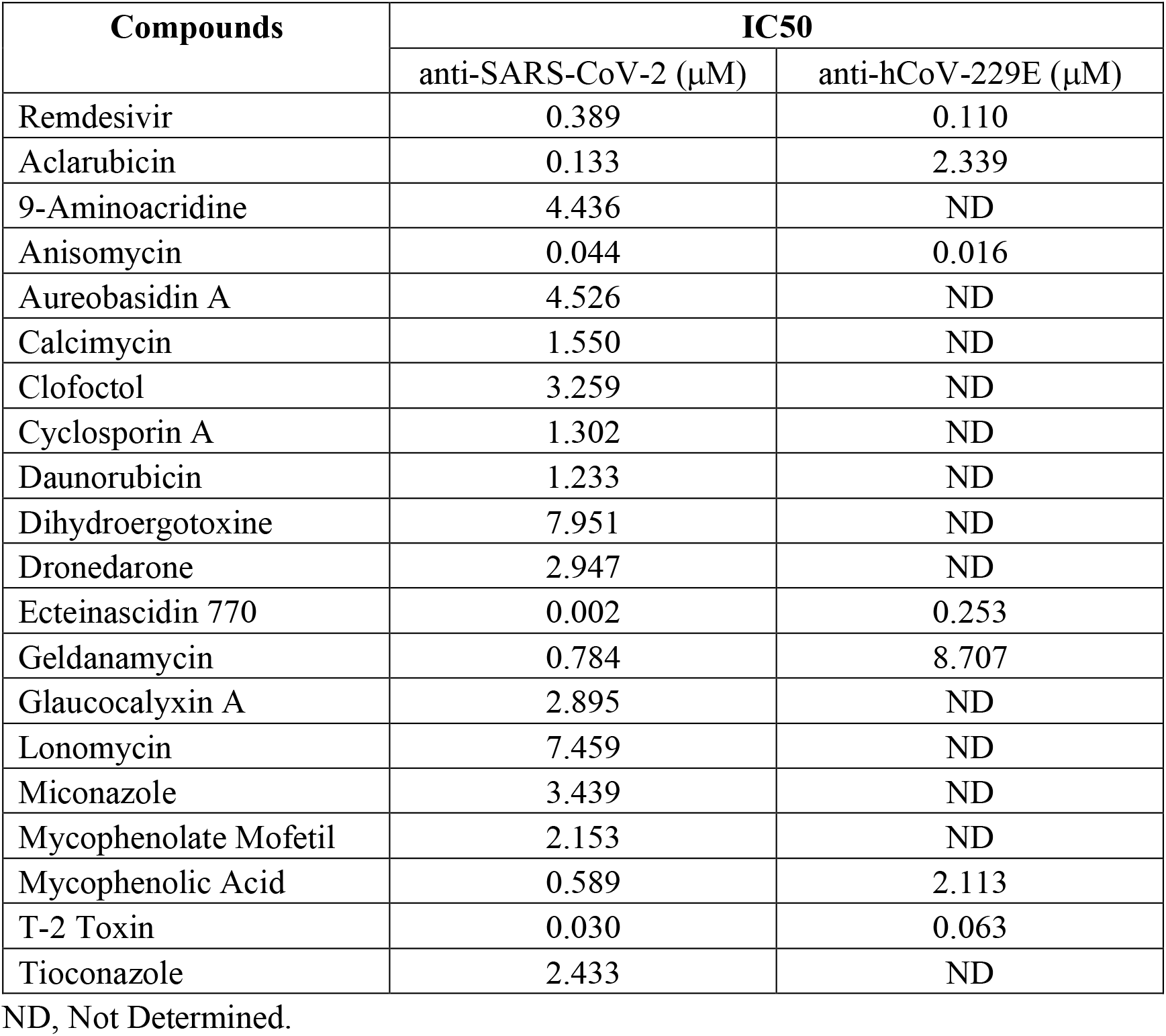
IC50 values of selected compounds inhibiting either SARS-CoV-2 or hCoV-229E.

### Aclarubicin and ET-770 inhibit SARS-CoV-2 RNA-dependent RNA polymerase

Aclarubicin and ET-770 are DNA intercalating agents (39). We thus tested whether they interfere with the RNA replication activity of SARS-CoV-2 RdRp (coded by viral NSP12 gene). We previously established a GLuc (Gaussia luciferase) reporter assay in which GLuc mRNA has both the 5’ and 3’ untranslated region (UTR) of SARS-CoV-2 RNA so that this GLuc mRNA can be replicated and amplified by SARS-CoV-2 RdRp together with viral proteins NSP7 and NSP8 (Fig. 4A). Remdesivir exhibits strong inhibition in this RdRp-dependent reporter assay (29). We transfected this GLuc reporter cell line with plasmid DNA expressing SARS-CoV-2 RdRp, NSP7 and NSP8, followed by treatment with increasing doses of aclarubicin and ET-770. Results of GLuc assay showed that both aclarubicin and ET-770 strongly inhibited RdRp-dependent expression of GLuc, with IC50 values of 0.383 μM for aclarubicin and 28 nM for ET-770 (Fig. 4B). Aclarubicin and ET-770 were shown to bind to SARS-CoV-2 main protease 3CL in the in silico molecular docking studies (40, 41). We therefore used aclarubicin and ET-770 to treat HEK293T cells that express SARS-CoV-2 3CLPro and a EGFP-Rluc fusion protein containing a target peptide of 3CLPro (Fig. 4C) (30). We included the 3CLPro inhibitor nirmatrelvir as a positive control which inhibited the cleavage of fusion protein EGFP-Rluc by 3CLPro (Fig. 4D). Neither aclarubicin nor ET-770 showed measurable inhibition of 3CLPro (Fig. 4D). Therefore, aclarubicin and ET-770 suppress SARS-CoV-2 infection at least partly by inhibiting viral RNA replication by RdRp.

**Figure 4.**
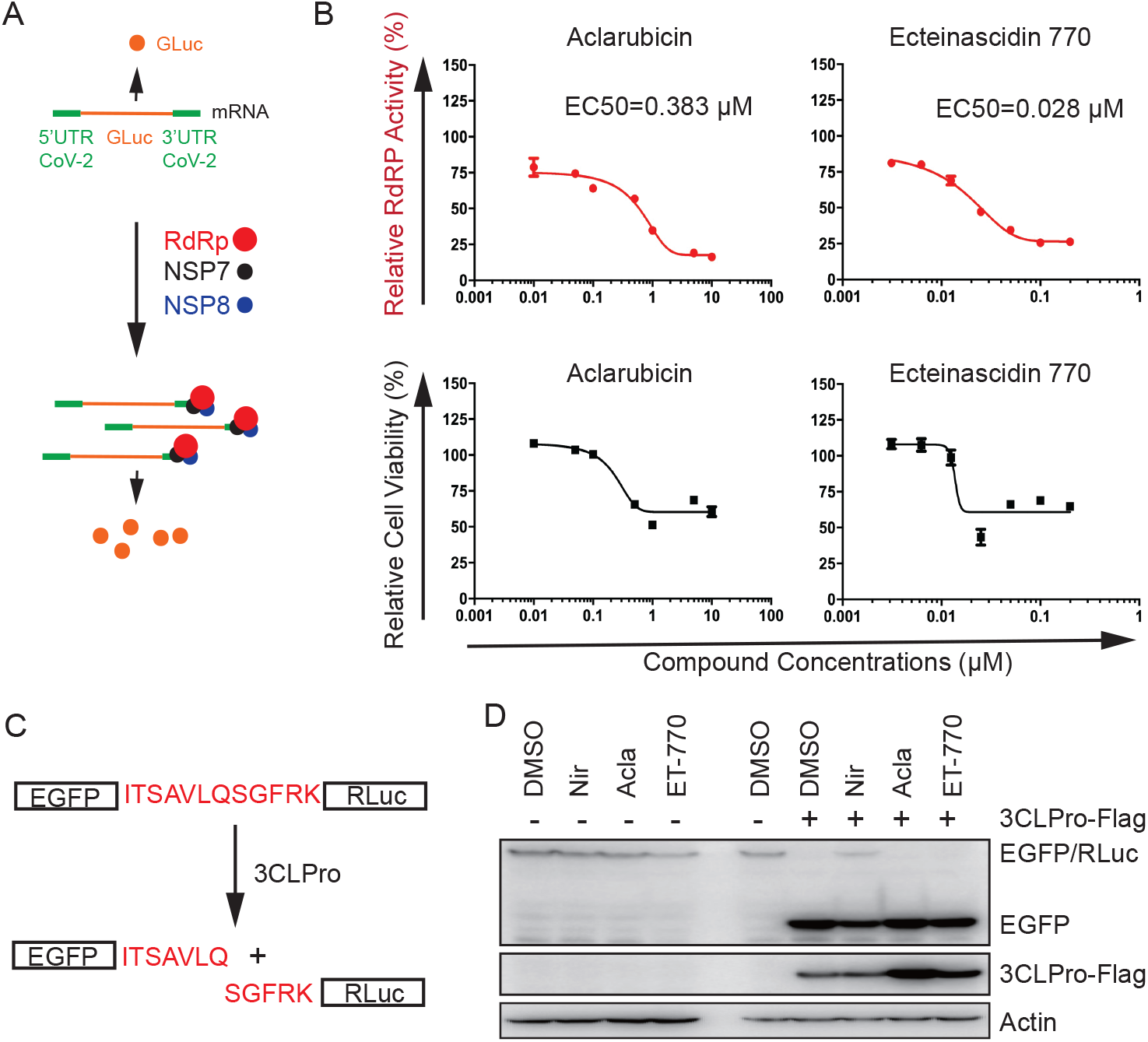
Aclarubicin and ecteinascidin 770 inhibit SARS-CoV-2 RdRp-mediated expression of GLuc. (A) Illustration of cell-based SARS-CoV-2 RdRp assay. The GLuc reporter mRNA has both the 5’ and 3’ UTRs of SARS-CoV-2 RNA which allow amplification by viral RNA polymerase complex NSP7/8/12 and produce higher levels of GLuc. (B) HEK293T cells expressing GLuc mRNA were transfected with NSP7, NSP8 and RdRp (NSP12), then treated with different concentrations of aclarubicin or ecteinascidin 770. EC50 value for each compound was calculated. Cytotoxicity of each compound in HEK293T cells was also measured. All drug concentrations were tested in triplicate. (C) Depiction of the EGFP-RLuc fusion protein bearing a linker sequence that can be cleaved by SARS-CoV-2 main protease 3CLPro. (D) HEK293T cells were transfected with plasmid DNA EGFP-Rluc and 3CLPro-Flag, then treated with nirmatrelvir (Nir, 1 μM), aclarubicin (Acl, 800 nM) or ecteinascidin 770 (ET-770, 10 nM) for 48 hours before cells were harvested and examined in Western blots using antibodies against EFP, Flag or actin. Results shown represent two independent transfection and drug treatment experiments.

### Anthracyclines inhibit SARS-CoV-2 infection

Aclarubicin is a member of the anthracycline drug family (42). Anthracyclines are commonly used to treat a variety of cancers such as leukemia, lymphoma, breast cancer, and lung cancer (43). Daunorubicin, doxorubicin, epirubicin and idarubicin are among the most frequently prescribed cancer drugs (Fig. 5A). We therefore tested whether these latter four anthracyclines also inhibit SARS-CoV-2 infection. Mitoxantrone is an anthraquinone, has similar molecular structure to daunorubicin, previously reported to inhibit SARS-CoV-2 entry (44, 45), and is thus tested as a control (Fig. 5A). The results showed that idarubicin inhibited SARS-CoV-2 to a similar degree of that by aclarubicin (Fig. 5B). It was reported that anthracyclines activate the expression of interferon stimulated genes (ISGs) to inhibit viral replication (46–48). We thus treated Calu-3 cells with these anthracyclines at 0.4 μM and 2 μM, and examined ISG expression by Western blotting. Indeed, daunorubicin, doxorubicin, epirubicin and idarubicin elevated the expression of MxB, IFITM3 and ISG56, with idarubicin showing the strongest activation (Fig. 5C). Mitoxantrone and aclarubicin did not activate these ISGs (Fig. 5C). The ISG activation activity of these anthracyclines was further confirmed by their stimulating the expression of luciferase that is controlled by the interferon-stimulated response element (ISRE) (Fig. 5D). We further measured levels of IFN-λ in the culture supernatants of treated Calu-3 cells. Daunorubicin, doxorubicin and idarubicin all significantly increased IFN-λ production compared to the DMSO control (Fig. 5E).

**Figure 5.**
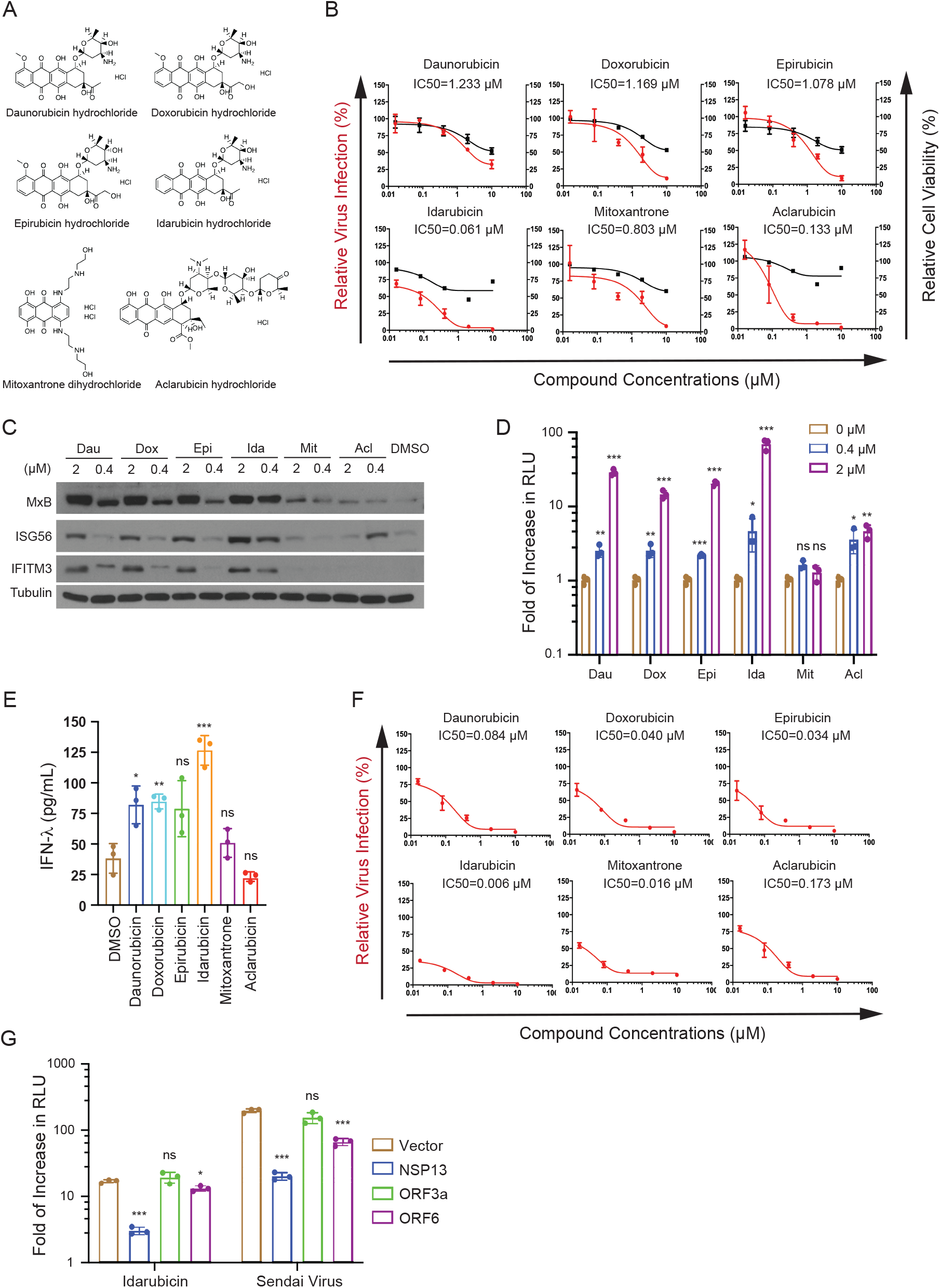
Inhibition of SARS-CoV-2 by anthracyclines. (A) Molecular structures of daunorubicin, doxorubicin, epirubicin, idarubicin, mitoxantrone and aclarubicin. (B) Effect of anthracyclines on CoV-2/NLuc infection when added at the time of infection. Virus infection was determined by measuring NLuc activity in the infected Calu-3 cells. Values of NLuc activity in the control infection (1% DMSO) are arbitrarily set as 100. All drug concentrations were tested in triplicate infections. (C) Effects of anthracyclines on the expression of MxB, ISG56 and IFITM3 in Calu-3 cells as examined in Western blots. Each drug was tested at 0.4 μM and 2 μM. Tubulin was detected as a control. Results shown represent two independent experiments. (D) The ISRE-Luc reporter plasmid DNA was transfected into HEK293T cells followed with treatment by anthracyclines at 0.4 μM and 2 μM. Luciferase activity was measured 48 hours after treatment. Luciferase values in cells exposed to 1% DMSO are arbitrarily set as 1. Each drug concentration was tested in triplicate transfections. (E) Levels of IFN-λ in culture supernatants of Calu-3 cells treated with anthracyclines at 2 μM in triplicate. (F) CoV-2/NLuc infection of Calu-3 cells that were pretreated with different concentrations of anthracyclines for 16 hours. NLuc values in the control infections (1% DMSO) are arbitrarily set as 100. (G) SARS-CoV-2 proteins NSP13 and ORF6 suppress the activation of ISRE-Luc reporter by idarubicin. HEK293T cells were either transfected with ISRE-Luc reporter plasmid DNA alone or together with NSP13, ORF3a or ORF6, followed with treatment of idarubicin (2 μM) or Sendai virus for 24 hours before luciferase activity was measured. The fold of change in luciferase expression activated by idarubicin or Sendai virus for each transfection condition was presented. All drug concentrations were tested in triplicate. p values are indicated as *p<0.05; **p<0.01; ***p<0.001; ns, not significant.

We reasoned that if idarubicin inhibits SARS-CoV-2 by activating IFN and ISGs, then pretreatment of Calu-3 cells with idarubicin should lead to stronger inhibition of SARS-CoV-2. Indeed, we observed a 10-fold lower IC50 associated with idarubicin pretreatment. In contrast, pretreatment with aclarubicin did not result in stronger inhibition of SARS-CoV-2 (Fig. 5F), which agrees with the inability of aclarubicin to activate ISGs in Calu-3 cells. Notably, pretreatment of Calu-3 cells with daunorubicin, doxorubicin or epirubicin also led to stronger inhibition of SARS-CoV-2 (Fig. 5F), which is consistent with their activation of ISGs and IFN-λ. Mitoxantrone, which did not activate ISG expression, inhibited SARS-CoV-2 to a higher fold when used to pretreat Calu-3 cells (Fig. 5F), which is likely because mitoxantrone targets the SARS-CoV-2 Spike-heparan sulfate complex and impairs virus entry (45).

SARS-CoV-2 encodes IFN antagonists that suppress IFN response and promote viral replication (49). We asked whether IFN stimulation by idarubicin can be countered by these SARS-CoV-2 proteins. We first examined the IFN antagonization activity of SARS-CoV-2 proteins NSP13, ORF3a and ORF6 which were previously reported to suppress IFN response by different mechanisms (50–53). In the experiment using the Sendai virus to activate ISRE-Luc which reports the degree of IFN activation, we observed that NSP13 and ORF6 suppressed activation of luciferase expression by Sendai virus infection (Fig. 5G). We then used idarubicin to treat cells that were either transfected with ISRE-Luc alone or co-transfected with SARS-CoV-2 genes. The results showed idarubicin increased luciferase expression by 15 folds, and this activation was effectively suppressed by viral NSP13 protein and to a lesser extent, by ORF6 (Fig. 5G). These data suggest that idarubicin may lose some of its antiviral effect when used to treat SARS-CoV-2-infected cells, because the expressed SARS-CoV-2 proteins NSP13 and ORF6 can suppress IFN activation by idarubicin. This explains why pretreatment of cells with idarubicin leads to a greater inhibition of SARS-CoV-2 infection, because pretreatment gives idarubicin time to stimulate ISG expression which establishes antiviral state before exposure to SARS-CoV-2 infection.

Together, these data demonstrate that different anthracyclines inhibit SARS-CoV-2 infection of Calu-3 cells to different degrees and by different mechanisms, with daunorubicin, doxorubicin, epirubicin and idarubicin stimulating the expression of ISGs, whereas aclarubicin suppresses viral RNA replication by viral RdRp.

## DISCUSSION

Through testing 620 microbial metabolites, we identified 6 compounds that inhibited SARS-CoV-2 with an IC50 lower than 1 μM. Some of these compounds have been shown by other groups to inhibit SARS-CoV-2. For example, anisomycin was reported to inhibit SARS-CoV-2 infection of Vero E6 cells at IC50 of 31.4 nM through stimulating IFN-β production (54), which is similar to the IC50 value of 44 nM measured in this study. Anisomycin has been shown to inhibit hCoV-OC43 (55) and hCoV-229E in this study, suggesting a pan-coronavirus activity of this compound. A second example is mycophenolic acid which is an inhibitor of inosine-5’-monophosphate dehydrogenase 2 (IMPDH2) and inhibits SARS-CoV-2 at least partially through suppressing viral protein NSP14-mediated activation of NF-κB (56, 57). We also observed the anti-SARS-CoV-2 activity of cyclosporin A which was evaluated in clinical trials as a potential treatment for COVID-19 (58).

It is not surprising to observe inhibition of SARS-CoV-2 by geldanamycin, given that a large number of viruses have been reported subject to geldanamycin inhibition. These include encephalomyocarditis virus (EMCV) (59), bombyx mori nucleopolyhedrovirus (60–62), pseudorabies virus (63), H5N1 influenza A virus (64), porcine circovirus 2 (65), chikungunya virus (66, 67), porcine reproductive and respiratory syndrome virus (68), enterovirus 71 (69, 70), herpes simplex virus type 2 (71), human cytomegalovirus (71, 72), Theiler’s murine encephalomyelitis virus (73), Ebola virus (74), hepatitis E virus (75), influenza virus (76), hepadnavirus (77), hepatitis B virus (77), flock house virus (78), vesicular stomatitis virus (79), hepatitis C virus (80, 81), herpes simplex virus type 1 (82, 83), vaccinia virus (84), RT and Rauscher leukemia virus (85). This broad antiviral activity of geldanamycin is primarily attributed to its antagonism of HSP90. Indeed, HSP90 is well known to assist and promote viral infection as a molecule chaperone (86). Similarly, other inhibitors of HSP90 were shown to inhibit SARS-CoV-2, including 17-AAG (87) and SNX-5422 (88), rendering HSP90 a promising target for the development of a pan antiviral drug (89).

We discovered the anti-SARS-CoV-2 activity of two anti-cancer drugs, aclarubicin and ecteinascidin 770. Aclarubicin was initially isolated from Streptomyces galilaeus in 1975 as an oligosaccharide anthracycline antibiotic, used to treat cancer by intercalating into DNA and acting as a topoisomerase II poison (90). We observed that aclarubicin inhibited SARS-CoV-2 and its variants in cultured cells with IC50 values in the range of 100 nM. Molecular docking analysis suggested aclarubicin as a potential inhibitor of SARS-CoV-2 main protease 3CL^pro^ (40); however, results of our cell-based assays did not reveal measurable inhibition of SARS-CoV-2 main protease by aclarubicin (Fig. 4D). Instead, SARS-CoV-2 RdRp was inhibited by aclarubicin in cell-based assays (Fig. 4B).

Aclarubicin does not share the mechanism of inhibiting SARS-CoV-2 by other anthracyclines such as doxorubicin and idarubicin which activate the expression of IFN and antiviral ISGs. This function of doxorubicin and idarubicin was previously reported by other groups (91), and confers inhibition of not only SARS-CoV-2 as we observed in this study, but also other viruses, including EMCV (91), Ebola virus (47), and hepatitis B virus (46). Viruses have evolved complex anti-IFN mechanisms, and some of the viral antagonists of IFN have been shown to be ineffective in suppressing IFN activation by doxorubicin, such as VP35 proteins of Ebola virus and Marburg virus, NS1 proteins of influenza A virus and respiratory syncytial virus, V and W proteins of Nipah virus, because these viral proteins target RIG-I, thus abrogate RIG-I sensing of viral RNA, whereas doxorubicin stimulates IFN and ISGs by activating IRF1 and/or IRF3 which are not targeted by these above viral proteins (47). In contrast, SARS-CoV-2 protein NSP13 effectively suppressed IFN activation by idarubicin. This is likely because NSP13 targets the late steps of IFN signaling pathway, such as activation of STAT1/STAT2 proteins (92), which are also required for idarubicin to increase ISG expression. Therefore, the effectiveness of idarubicin as a treatment for viral infection such as SARS-CoV-2 can be compromised by the multi-layered viral antagonization of IFN response.

In summary, we have identified several microbial metabolites with promising anti-SARS-CoV-2 activity, including aclarubicin that strongly inhibits SARS-CoV-2, and is worth further exploring as a potential SARS-CoV-2 inhibitor.

## ACKNOWLEDGEMENTS

We would like to thank Dr. Irah King for insightful discussions of microbial metabolites, Dr. Matthias Gotte, Egor Tchesnokov, and Emma Woolner for the help with SARS-CoV-2 RdRp assay. The authors thank the Laboratoire de Santé Publique du Québec for the authentic Omicron BA.1 virus. This study is supported by CIHR operating grant OV4-170641 to C.L., Sentinelle COVID Quebec, Jewish General Hospital Foundation, McGill Interdisciplinary Initiative in Infection and Immunity (MI4), and the biosafety level 3 facility at the Research Institute of the McGill University Health Centre. This work was also partially supported by a CIHR operating Pandemic and Health Emergencies Research grant #177958 to A.F. A.F. is the recipient of Canada Research Chair on Retroviral Entry no. RCHS0235 950-232424.

